# Selective targeting of RalA with intrabodies impairs Triple Negative Breast Cancer metastasis

**DOI:** 10.64898/2026.01.09.698394

**Authors:** Charlotte Sarre, Katerina Jerabkova-Roda, Vincent Denis, Giulia Bertolin, Marc Tramier, Louis Bochler, Cristina Liboni, Ignacio Busnelli, Quentin Frenger, Annabel Larnicol, Frédéric Gros, Olivier Lefebvre, Laurence Guglielmi, Jacky G. Goetz, Vincent Hyenne, Pierre Martineau

**Author notes:** Equal contribution (Charlotte Sarre, Katerina Jerabkova-Roda). These authors share senior authorship (Jacky G. Goetz, Vincent Hyenne, Pierre Martineau).

## Abstract

This study successfully developed and validated isoform-specific intrabodies targeting the highly homologous Ral oncoproteins, key effectors in cancer progression. Phage display was used to isolate single-chain variable fragment (scFv) clones that recognize specifically RalA (C1-A, G5-A), RalB (F6-B), or both paralogs (A12-AB). Using lentiviral transduction, these intrabodies were stably expressed as GFP fusions in murine breast cancer 4T1 cells. The anti-RalA clones C1-A and A12-AB demonstrated clear colocalization with RalA, confirming their binding activity inside the cell. We further confirmed their activity in cells by analyzing Ral-dependent pathways. All intrabodies targeting RalA (C1-A, G5-A, A12-AB) but not the one specific to RalB inhibited mitochondrial fission, a known RalA function. All clones, except the pan-Ral one, altered the endo-lysosome pathway by decreasing lysosome number. Furthermore, the RalA-specific C1-A clone reduced lysosomes size, and uniquely and strongly reduced extracellular vesicle secretion, highlighting its distinct inhibitory potential. In an orthotopic Triple-Negative Breast Cancer (TNBC) mouse model, The C1-A RalA specific clone significantly but weakly reduced primary tumor growth, but exerted a powerful anti-metastatic effect, dramatically reducing lung metastases, with a complete abolition of metastases observed in 2/5 mice.

In summary, these potent, isoform-specific Ral intrabodies act as effective intracellular inhibitors, successfully modulating RalA-specific functions in cells and offering a promising therapeutic strategy for significantly suppressing tumor growth and metastasis in vivo.

## Introduction

Intrabodies are functional intracellular antibodies emerging as powerful tools for drug discovery through their ability to target intracellular proteins. Today, phage display technologies allow the generation of highly selective intrabodies (Hsiue *et al*, 2021). Proteins difficult to inhibit with traditional small molecules become now accessible despite their flat surfaces (Douglass *et al*, 2021). Unlike classical silencing approaches and inhibitor compounds that shut down the full pool of target proteins in cell, intrabodies have the ability to be more selective (Cattaneo & Chirichella, 2019). Indeed, intrabodies can be selected to bind specific conformational states, misfolded proteins, post-translational modifications or even isoforms with high homology, allowing them to regulate signaling pathways through a variety of mechanisms (De Groof *et al*, 2021). Among them, intrabodies can prevent protein-protein interactions by interfering with complex assembly and effector recruitment, they can perturb protein activation via post translational modification, mark the protein for degradation or perturb its subcellular localization (Lin *et al*, 2020). These make intrabodies powerful tools for blocking dysfunctional or hyperactive signaling pathways in diseases such as cancer where aberrant signaling drives disease progression.

We took advantage of the intrabody technology to target RalA small GTPases, a protein that switches between a GDP-bound inactive state and a GTP-bound active state in cells. This protein constitutes ideal yet challenging targets. RalA protein and its RalB paralog are critical downstream effectors of Ras, whose activity and expression are increased in a large number of cancers, controlling various steps of tumor development from tumor initiation to tumor growth and metastatic progression (Yan & Theodorescu, 2018). However, despite showing a high degree of homology (reaching 82% of similarity in human), RalA and RalB can have redundant, specific or antagonistic functions depending on the cancer type (Richardson *et al*, 2022). In different models of triple negative breast cancer for instance, RalA and RalB were shown to have opposite effects on primary tumor growth and metastasis (Thies *et al*, 2021) (Ghoroghi *et al*, 2021). By contrast, in other cancer types, such as lung carcinoma, RalA and RalB have complementary or redundant roles (Peschard *et al*, 2012; Guin *et al*, 2013). These differences could be explained by the fact that RalA and RalB are pleiotropic proteins controlling various essential cellular processes through a number of effectors, including endosomal trafficking, cytoskeleton remodeling, mitochondria fission, nutrient sensing or exosome secretion (Richardson et al., 2022). Some of these functions, such as late endolysosome maturation and exosome secretion, involve both RalA and RalB (Ghoroghi *et al*, 2021), while others are paralog specific, such as mitochondria fission, which is only controlled by RalA (Kashatus *et al*, 2011). Consequently, RalA and RalB affect multiple cellular phenotypes, including cell migration and invasion, cell proliferation, cell metabolism in both healthy and pathological contexts (Apken & Oeckinghaus, 2021).

Given their importance in various cancer, several Ral inhibitors have been developed over the past years. Yet, similar to Ras, Ral proteins are challenging pharmaceutical targets, due to the lack of deep hydrophobic pockets and high GTP affinity. Several approaches have been developed, including inhibition of Ral relocalization using a geranylgeranyl (GG) inhibitor (Falsetti *et al*, 2007; Yan *et al*, 2014) or design of stapled-peptides to selectively bind GTP-bound RalA and RalB (Thomas *et al*, 2016; Hurd *et al*, 2021). Two small molecule inhibitors, RBC8 and its derivate BQU57, were shown to stabilize Ral proteins in their inactive forms (Yan et al., 2014) and displayed initial promising results in preclinical in vivo models, but the identification of off-targets effects questioned their specificity (Walsh *et al*, 2019; Han *et al*, 2024). More recently, a new small-molecule inhibitor, OSURALi, inhibiting both RalA and RalB GTP binding, showed a Ral-dependent specific cytotoxicity in TNBC cell lines (Han *et al*, 2024). Yet, its potency has not been evaluated in vivo. To date, no paralog specific Ral inhibitor has been developed and there is no Ral inhibitor in clinical use.

Starting from a highly diverse hyper-stable synthetic scFv library, we have generated anti-RalA intrabodies able to bind both paralogs or specifically RalA. We found that these new RalA inhibitors are expressed and functional in a carcinoma cell line where they inhibit specific RalA functions. Importantly, one of them almost fully inhibits lung metastasis in a murine model of triple negative breast cancer, despite a minimal effect on primary tumor. This work highlights the utility of intrabodies for deciphering complex and interconnected pathways, and it paves the way for new therapeutic options, based on future small-molecule inhibitors that mimic intrabody effects (Villoutreix *et al*, 2011; Mazuc *et al*, 2008; Quevedo *et al*, 2018).

## Results

### Screening of a library of scFv identifies RalA-specific intrabodies

To generate antibodies recognizing Ral GTPases, we screened the HUSCI2 library of humanized synthetic scFv antibody by phage display (Fig. 1A) (Abba Moussa *et al*, 2025). This library is a mono-framework library based on clone scFv13R4, a scFv selected for its high stability (Martineau & Betton, 1999) and cytoplasmic expression in E. coli and in mammalian cells (Martineau *et al*, 1998; Sibler *et al*, 2003). Diversity in this mono-framework library was introduced at 16-21 positions chosen in the six complementarity-determining regions. Clones were isolated by affinity selection (panning) in three selection campaigns (Fig. 1A). First, the naive library was panned on purified RalA protein, yielding essentially clones recognizing both RalA and RalB and a single RalA-specific clone, due to high sequence homology of both paralogs. Second, to obtain RalA specific clones, a selection was performed with an initial depletion on RalB, followed by affinity selection on RalA. Third and conversely, to obtain RalB specific clones, the library was depleted on RalA, followed by selection on RalB. In total, after sequencing we obtained 20 different pan-Ral, 5 RalA and 5 RalB specific clones and verified their binding specificity by ELISA (Fig. 1B). The 5 anti-RalA scFv clustered in 3 families (based on VH-CDR3 sequences; Supplementary Fig. S1) and one clone of each group (C1-A, G5-A and D10-A) was cloned as an eGFP fusion in a lentiviral vector and tested for expression in HeLa cells by cytometry and live microscopy (Supplementary Fig. S2). Clone D10-A showed strong fluorescent spots, a clear indication of aggregation in the cell, and it was discarded. On the opposite, clones C1-A and G5-A showed a homogenous fluorescence with no visible aggregates and were retained for the rest of the study. As controls we retained one RalB-specific clone, F6-B, and one pan-Ral scFv equally reacting with both paralogs (A12-AB) (Fig. 1B & Supplementary Fig. S1).

**Figure 1.**
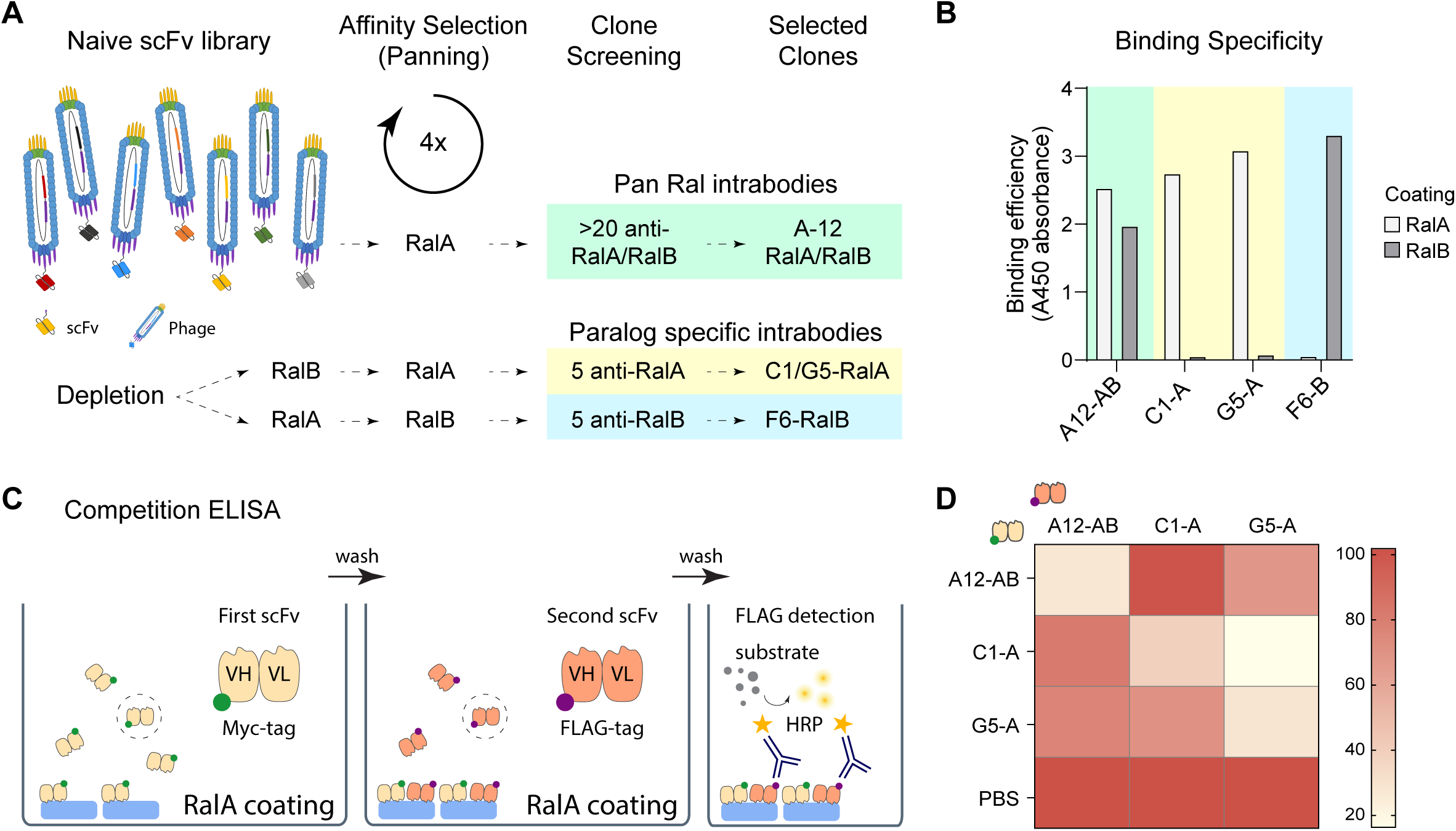
Selection and characterization of RalA specific intrabodies. **A**) Schematic representation of intrabodies selection by panning on immobilized Ral, using the scFv library. Paralog-specific clones are selected in an additional step by depletion on one Ral paralog followed by selection on the other paralog. Green= pan-specific, yellow= RalA-specific, blue= RalB-specific clones. **B**) Binding specificity of selected clones, determined by ELISA (coating: white= RalA, gray= RalB). Green= pan-specific, yellow= RalA-specific, blue= RalB-specific clone. ScFv concentration: 12 μg/mL. **C**) Principle of competition ELISA and detection. **D**) Heatmap representing competition ELISA between the 3 anti-RalA clones. White = high competition, clones recognize the same site, red = low competition, clones do not bind to the same site.

To determine if the different selected scFvs bind overlapping epitopes on Ral GTPases we performed competitive ELISA. Purified Ral proteins were coated on microtiter plates and a saturating concentration of each scFv (myc tagged) was independently added to block the recognized epitope, before adding another scFv with a different tag (FLAG; Fig. 1C). We observed that the 3 scFvs recognizing anti-RalA, C1-A, G5-A and A12-AB, partially but not fully inhibited each other, suggesting that these 3 antibodies recognize close epitopes, different for A12 and C1 (Fig. 1D & Supplementary Fig. S3A). Because of the size of scFvs (30 kDa) in comparison to RalA (23 kDa), partial inhibition even with non-identical binding sites was expected. Similarly, F6-B and A12-AB clones did not inhibit each other and recognize distant and different epitopes on RalB protein (Supplementary Fig. S3A).

### The expression of Ral-specific intrabodies is inducible and stable in a TNBC model

To generate functional intrabodies (intracellular antibodies), we stably expressed, using lentiviral infection, the anti-Ral scFvs fused to eGFP under the control of TetOn inducible promoter in murine TNBC 4T1 cells. While this model allows to probe tumor progression of TNBC in a orthotopic context, we had also previously shown that Ral proteins control endo-lysosome maturation and metastasis formation (Ghoroghi *et al*, 2021). As a negative control, we used non-targeting scFv directed against *E. coli* β-galactosidase. Because this control clone was used to construct the scFv library, it shares around 94% of identity (93.6 -- 95.1%) with the anti-Ral scFvs.

We demonstrated previously that misfolded and aggregated scFv are quickly degraded by the cell proteasome and that the intrabody expression and solubility directly scales with GFP expression (Guglielmi *et al*, 2011). To assess the intrabody stability in 4T1 cells over time, we monitored the scFv-GFP expression by flow cytometry and observed robust GFP signal 6h after doxycycline (Dox) treatment, lasting up to 48 h (Fig. 2A & Supplementary Fig. S3B). At the peak of expression, the signal showed a 7.8 to 21-fold increase over background (respectively for C1-A and A12-AB) and a high proportion of positive cells (72-93%; Fig. 2B). The mean GFP fluorescence intensity at 24h post-induction was similar for all clones (MFI: 2225 – 3906), except for the RalA-specific clone C1-A, which showed a 2- to 3-fold lower GFP signal (MFI: 1144). Since intrabodies can face misfolding and aggregation upon expression (Guglielmi *et al*, 2011), we assessed their solubility and subcellular distribution using live fluorescent imaging. All selected clones showed homogenous GFP distribution with no visible aggregates (Fig. 2C). To confirm these results, we analyzed the expression of the scFv-GFP clones by western blot using an anti-GFP antibody. As shown in figure 2D, all cells expressed the complete scFv-GFP fusion protein at the anticipated molecular weight of 55 kDa, but at different levels of expression, confirming the cytometry analysis (Fig. 2B). All the clones also presented additional bands corresponding to a proteolytic degradation in the linkers of the fusion (Fig. 2E). Non-T, A12-AB and F6-B were cleaved between the two domains of the scFv, generating a band at 41 kDa, corresponding to the VL-GFP fusion. The two anti-RalA specific clones, C1-A and G5-A, were cleaved between the scFv and the GFP, generating a band at 29.5 kDa, corresponding to free GFP with additional C-terminal tags (c-myc and His6). In particular, the control anti-RalB F6-B was mainly degraded and only a faint band corresponding to the full construct was detected. This does not prevent activity since the Fv can remain associated even after the cleavage of the scFv linker. Therefore, while all clones are properly expressed and can be used for functional studies, only some of them can be used for reliable GFP-based tracking of their intracellular location. Taken together, these results show that selected anti-Ral scFv, despite some instability and degradation, are expressed in the cytoplasm of mammalian cells.

**Figure 2.**
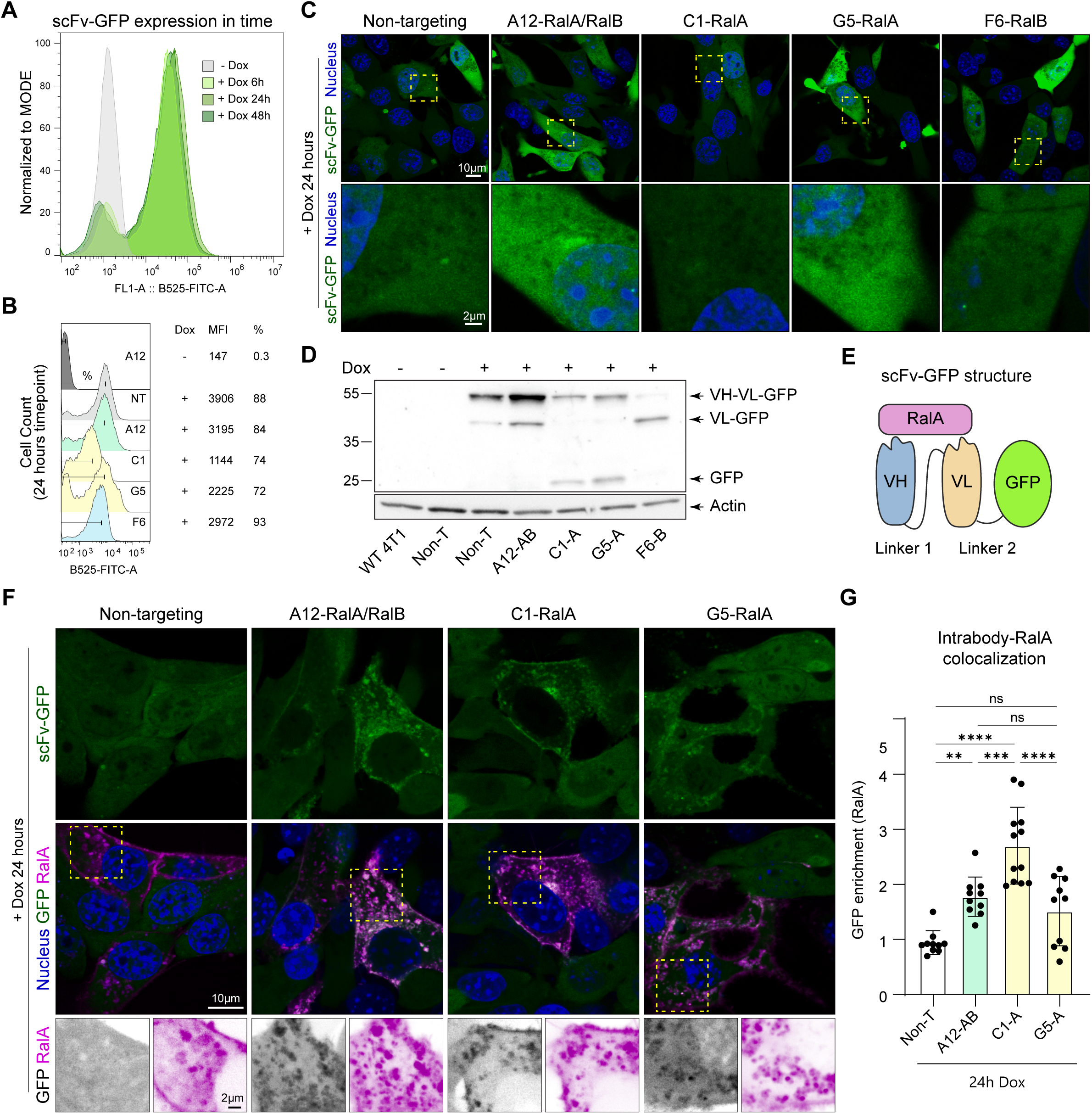
4T1 cells stably expressing anti-Ral intrabodies. **A**) Kinetics of GFP expression in 4T1 cells upon doxycycline treatment (0.5 µg/ml) analyzed by Flow cytometry at three different time points: 6 h, 24 h and 48 h after treatment (representative example, A12-AB clone). **B**) Flow cytometry analysis of GFP expression 24 h after doxycycline treatment in 4T1 cells showing mean fluorescence intensity (MFI) and the percentage of GFP-positive cells (%) for each clone. **C**) Intrabody-GFP subcellular distribution in 4T1 cells. Live-cell imaging by Spinning-disk confocal microscopy at 24 h after doxycycline treatment. Scale bar = 10 µm, zoom = 2 µm. **D**) Western blot analysis of intrabody-GFP expression in 4T1 stable cell lines 24 h upon doxycycline treatment (anti-GFP). **E**) Intrabody structure showing the VH, VL, GFP and the linkers. **F**) Representative confocal images of 4T1 cells transfected with tdTomato-RalA, stably expressing GFP-coupled intrabodies. Live cell imaging, as in C. Scale bar: 10 µm, zoom = 2 µm. Intensity adjusted to best visualize the localization patterns. **G**) Quantification of images in F: recruitment of GFP-intrabody to RalA-tdTomato shown as signal ratio (GFP colocalizing with RalA over total GFP). Values > 1 demonstrate intrabody recruitment. Mean ± SD = 0.941 ± 0.217, 1.773 ± 0.357, 2.704 ± 0.698, 1.515 ± 0.631. One dot represents one cell, n = 10 cells per condition in two independent experiments. Ordinary one-way ANOVA with Turkey’s multiple comparisons test, * p□<□0.05; ** p□<□0.01; *** p□<□0.001; **** p□<□0.0001.

We then wondered whether increasing the expression of RalA GTPases would be sufficient to concentrate the intrabodies on RalA-specific subcellular locations. We thus overexpressed tdTomato-RalA in 4T1 cells expressing RalA intrabodies (Fig. 2F & 2G). In line with our previous observations (Ghoroghi *et al*, 2021) and independently of which clone is co-expressed, we found that tdTomato-RalA localizes to the plasma membrane and to late endo-lysosomal compartments, which are mostly positive for lysotracker (Fig. 2F & Supplementary S4A). The non-targeting intrabody showed a homogenous diffused pattern, similar to the one observed where RalA is not over-expressed (Fig. 2C). In contrast, we observed that RalA-targeting GFP-intrabodies colocalize with overexpressed RalA both at the plasma membrane and on RalA positive endolysosomes (Fig 2F & 2G). In this context, both clone A12-AB and clone C1-A exhibit equally effective colocalization with RalA, despite the lower expression level of the latter (1.77 ± 0.11 and 2.7 ± 0.2, respectively) compared to the non-targeting clone (0.94 ± 0.07). The G5-A clone showed some but not significant colocalization, as expected since most of the GFP, because of the proteolytic cleavage, is no more linked with the intrabody. In conclusion, A12-AB and C1-A clone demonstrated binding activity in cells and can be used as intrabodies to modulate RalA activities.

### RalA intrabodies alter the endo-lysosomal system

Prompted by the successful creation of anti-RalA intrabodies in 4T1 cells, we next aimed at determining their efficiency as Ral inhibitors. We focused on Ral functions related to late-endosomal and lysosomal trafficking as well as secretion of Extracellular Vesicles (EVs) (Fig. 3A), since i) we have shown that Ral control endolysosome maturation and secretion of EVs (Hyenne *et al*, 2015; Ghoroghi *et al*, 2021) and ii) anti-RalA scFv colocalize to endo-lysosomes upon RalA overexpression (Fig. 2F and Supplementary Fig. S2). To do so, we assessed the impact of Ral intrabodies on lysosomes by analyzing their number and size in 4T1 cells by confocal live cell imaging (Fig. 3B). We found that three out of the four Ral Intrabodies (C1-A, G5-A and F6-B) significantly decreased the number of lysosomes per cell (Fig. 3C left). This decrease in lysosome number is coupled with an increase in their size for the C1-A and F6-B clones (Fig. 3C right). We further isolated EVs released by intrabody-expressing 4T1 cells by ultracentrifugation, identified their content by western blot and measured their concentration by nanoparticle-tracking analysis (Fig. 3D and Supplementary Fig. S4B). We found that expression of the C1-A clone is the only clone to induce a strong decrease in EV secretion in 4T1 cells. Altogether, those results show that most but not all Ral-targeting intrabodies alter the endo-lysosome pathway, regardless of the paralog targeted, but that the RalA-specific clone C1-A additionally blocked EV secretion.

**Figure 3.**
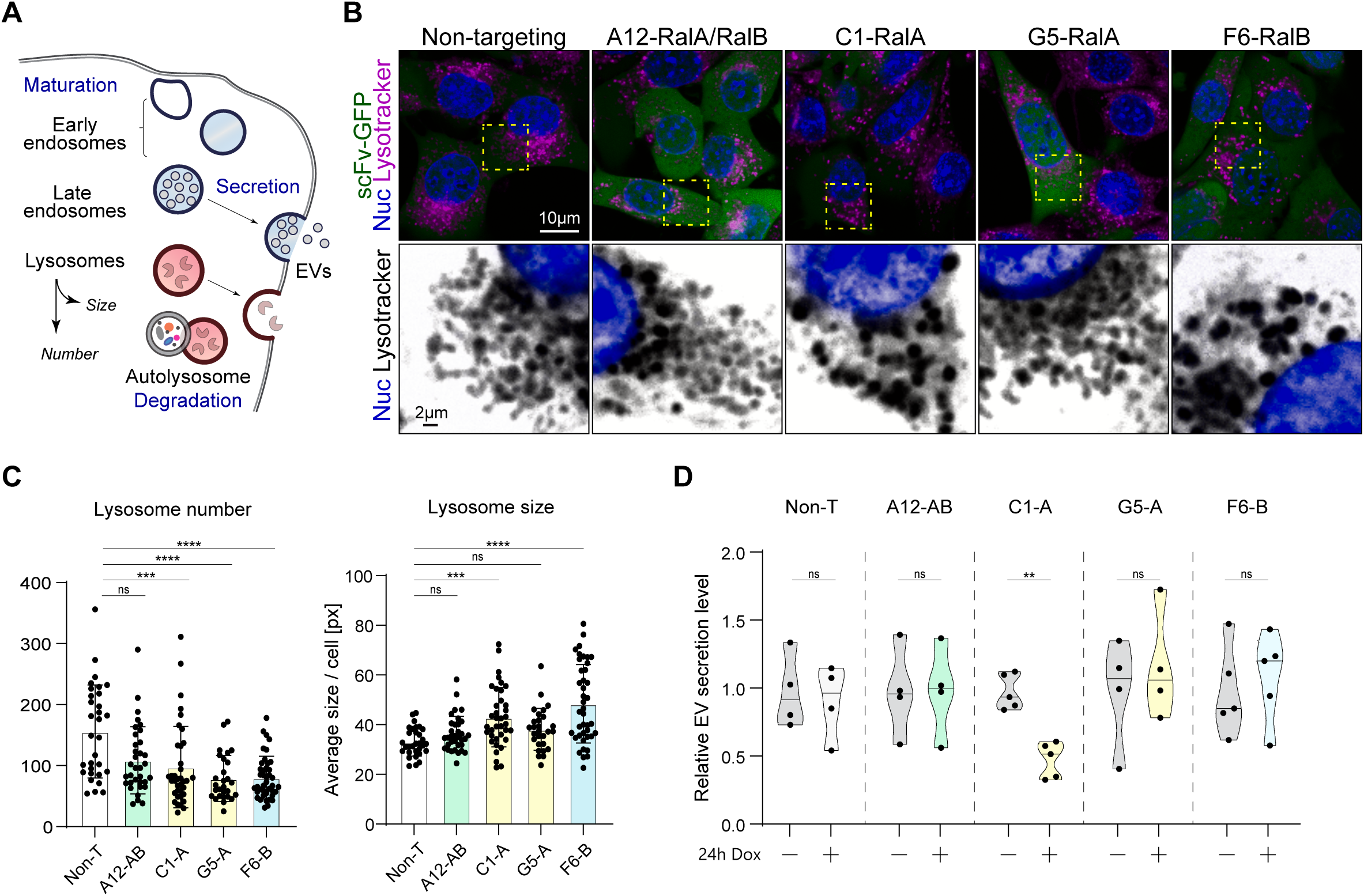
Intrabodies targeting RalA and RalB differentially modulate lysosome size and number, and extracellular vesicles secretion. **A**) Endo-lysosomal system. **B**) Representative confocal images of 4T1 cells. Live-cell spinning-disk imaging 24 h after doxycycline treatment. Blue = nucleus, green = intrabody-GFP, magenta = lysotracker. Scale bar = 10 µm, zoom = 2 µm. Intensity adjusted to visualize the morphology patterns. **C**) Quantification of B. Non-targeting (Non-T; n = 31), A12-AB (n = 34), C1-A (n = 37), G5-A (n = 28), F6-B (n = 80) cells in three independent experiments. Left: Lysosome number (Mean ± SD) = 155.9 ± 76.21, 108.9 ± 54.99, 97.46 ± 66.59, 78.89 ± 37.83, 80.27 ± 35.10. Right: Lysosome size (Mean ± SD) = 33.02 ± 5.643, 36.29 ± 7.103, 42.98 ± 11.97, 38.11 ± 8.407, 48.40 ± 15.79. Kruskal-Wallis with Dunn’s Multiple Comparison Test. One dot represents one cell. **D**) Quantification of extracellular vesicles (EV) numbers (Nanoparticle tracking analysis, NTA) isolated by ultracentrifugation (100,000 g pellet) from the supernatant of 4T1 cells induced or not for 24 h with doxycycline. Number of independent experiments for each clone: Non-targeting (Non-T; n = 4), A12-AB (n = 4), C1-A (n = 5), G5-A (n = 4), F6-B (n = 6). EV number (Mean ± SD) = 1.0 ± 0.27, 0.93 ± 0.27, 1.0 ± 0.33, 1.01 ± 0.33, 1.0 ± 0.12, 0.50 ± 0.12, 1.0 ± 0.0.40, 1.18 ± 0.40, 1.0 ± 0.33, 1.10 ± 0.33, normalized to mean of respective controls. Mann-Whitney test, two-tailed. * p□<□0.05; ** p□<□0.01; *** p□<□0.001; **** p□<□0.0001.

### RalA intrabodies inhibit mitochondria fission, a paralog specific function

We then wondered whether selected Ral intrabodies could inhibit RalA-specific functions. To address this question, we chose to study the impact of Ral intrabodies on mitochondria fission (Kashatus *et al*, 2011). Mitochondria were labelled with mitotracker and observed by live cell confocal imaging upon intrabodies expression (Fig. 4A). We observed that the three intrabodies targeting RalA (A12-AB, C1-A and G5-A) induced a significant increase in the mitochondrial aspect ratio, suggesting an impaired mitochondria fission. In contrast, the mitochondrial aspect ratio was not modified by non-targeting and RalB-specific (F6-B) clones (Fig. 4B).

**Figure 4.**
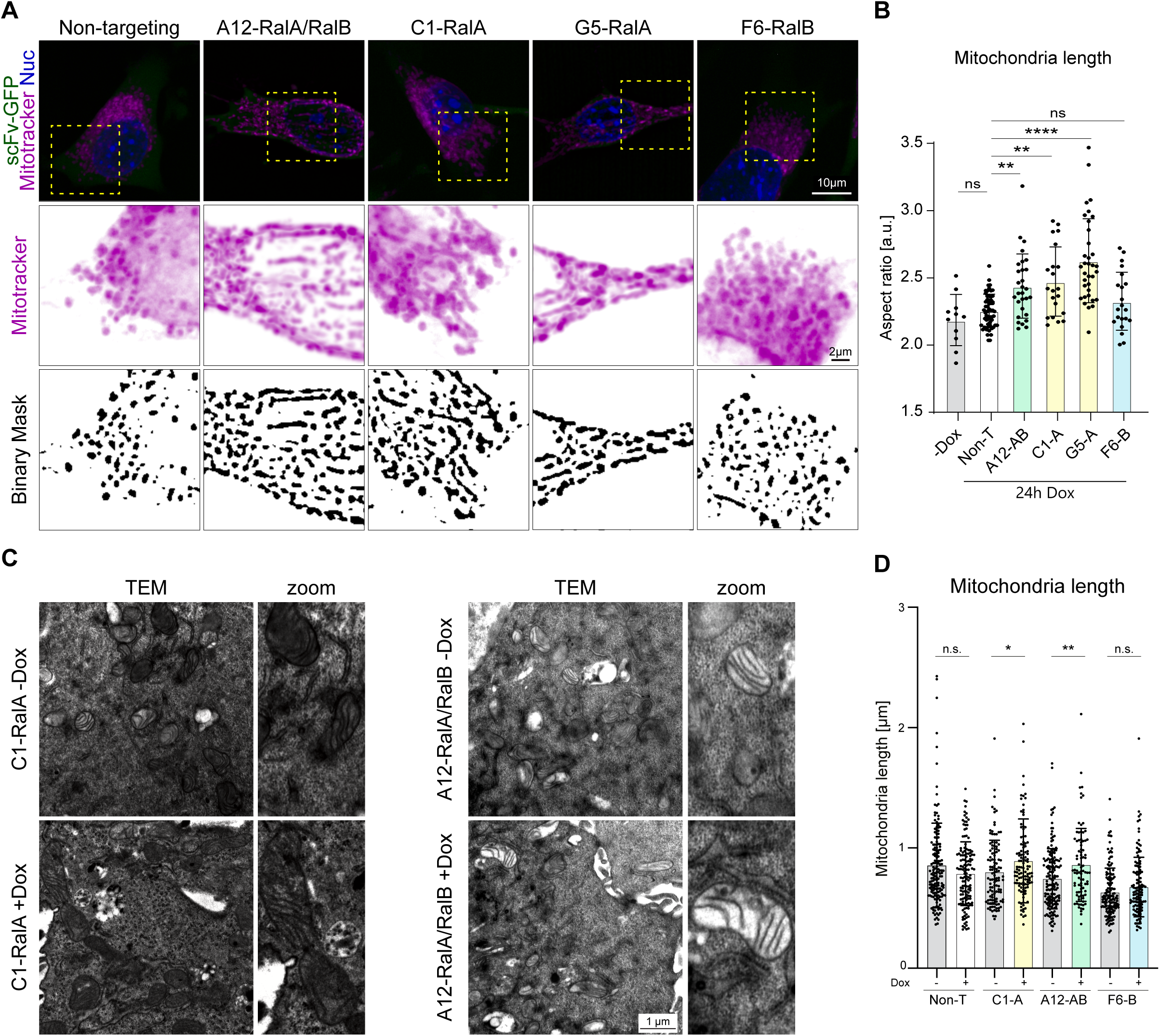
Intrabodies selectively inhibit RalA function in 4T1 cells. **A**) Mitochondrial network in living cells after 24 h of Dox induction, illustrating the mitochondrial network morphology for each clone. Representative confocal images of 4T1 cells stably expressing GFP-intrabodies. Live-cell spinning-disk imaging 24 h after doxycycline treatment. Blue = nucleus, green = scFv-GFP, magenta = mitotracker. Scale bar = 10 µm, zoom = 2 µm. **B**) Quantification of images in A showing mitochondria elongation (aspect ratio) for each clone. Non-T -Dox (n = 11); +Dox: Non-T (n = 56), A12-AB (n = 29), C1-A (n = 21), G5-A (n = 35), F6 (n = 21) cells. Aspect ratio (Mean ± SD) = 2.186 ± 0.1901, 2.257 ± 0.1226, 2.439 ± 0.2390, 2.473 ± 0.2571, 2.628 ± 0.3124, 2.326 ± 0.2155. Kruskal-Wallis with Dunn’s Multiple Comparison Test. One dot represents one cell, three independent experiments. **C**) Representative electron microscopy images of 4T1 cells expressing C1-A and A12-AB clones, with or without 24 h doxycycline treatment. **D**) Quantification of C. Number of mitochondria quantified: Non-T -Dox (n = 159), Non-T +Dox (n = 138), C1-A -Dox (n = 115), C1-A +Dox (n = 115), A12-AB -Dox (n = 162), A12-AB +Dox (n = 87), F6-B -Dox (n = 142), F6-B +Dox (n = 126). Mitochondria length (Mean ± SD) = 854.2 ± 351.2, 786.1 ± 264.8, 798.9 ± 265.6, 892.2 ± 348.0, 743.8 ± 249.6, 859.7 ± 303.0, 630.1 ± 202.2, 675.1 ± 247.0 nanometers. Mann-Whitney test, two-tailed. * p□<□0.05; ** p□<□0.01; *** p□<□0.001; **** p□<□0.0001.

To confirm these results and assess the mitochondria morphology at the ultrastructural level, we used transmission electron microscopy (Fig. 4C). Similarly, to what we observed with live imaging, the expression of clones targeting RalA induced a significant increase of mitochondrial size compared to untreated cells. In contrast, the expression of non-targeting clone and RalB-specific clone F6-B had no significant impact on mitochondrial length (Fig. 4D). These results collectively confirm that intrabodies targeting the RalA isoform effectively inhibit RalA-mediated mitochondrial fission.

### RalA and RalB specific intrabodies inhibit tumor growth and metastasis

Having demonstrated that Ral intrabodies are potent Ral inhibitors *in vitro,* we aimed at assessing their potential in controlling Ral-dependent tumor progression *in vivo*. Exploiting 4T1 cells in a orthotopic model which allows to concomitantly probe primary mammary tumor growth and lung metastasis formation, we had previously shown that inhibiting either RalA or RalB play on cancer progression (Ghoroghi *et al*, 2021). To test the efficacy of Ral intrabodies *in vivo*, we performed an orthotopic injection of four clones (RalA specific C1-A and controls RalB specific F6-B, pan-specific RalA/RalB A12-AB and non-targeting) and followed the tumor growth over time (Fig. 5A). We were able to detect expression of the intrabodies in all tumors after 33 days, demonstrating an efficient induction and a strong stability *in vivo*, despite variability in the expression level between clones and mice (Fig. 5B). Of note, regarding the anti-RalB clone, the full F6-B-GFP fusion was unstable and cleaved by the cell, as also observed *in vitro* (Fig. 2D), but the full-length fusion was however present, except in mouse m2 (Fig. 5B). When analyzing primary tumor growth, we found that tumors expressing the C1-A, F6-B or A12-AB anti-Ral intrabodies grew at a reduced pace compared to control tumors, resulting in a significant reduction in overall tumor volume at day 33 (Fig. 5C left). Tumors expressing the anti-RalB F6-B clone displayed the most important delay in tumor growth and were the only group to show a significant reduction in tumor weight at the end of the experiment, despite the low stability of this intrabody-GFP fusion (Fig. 5C right).

**Figure 5:**
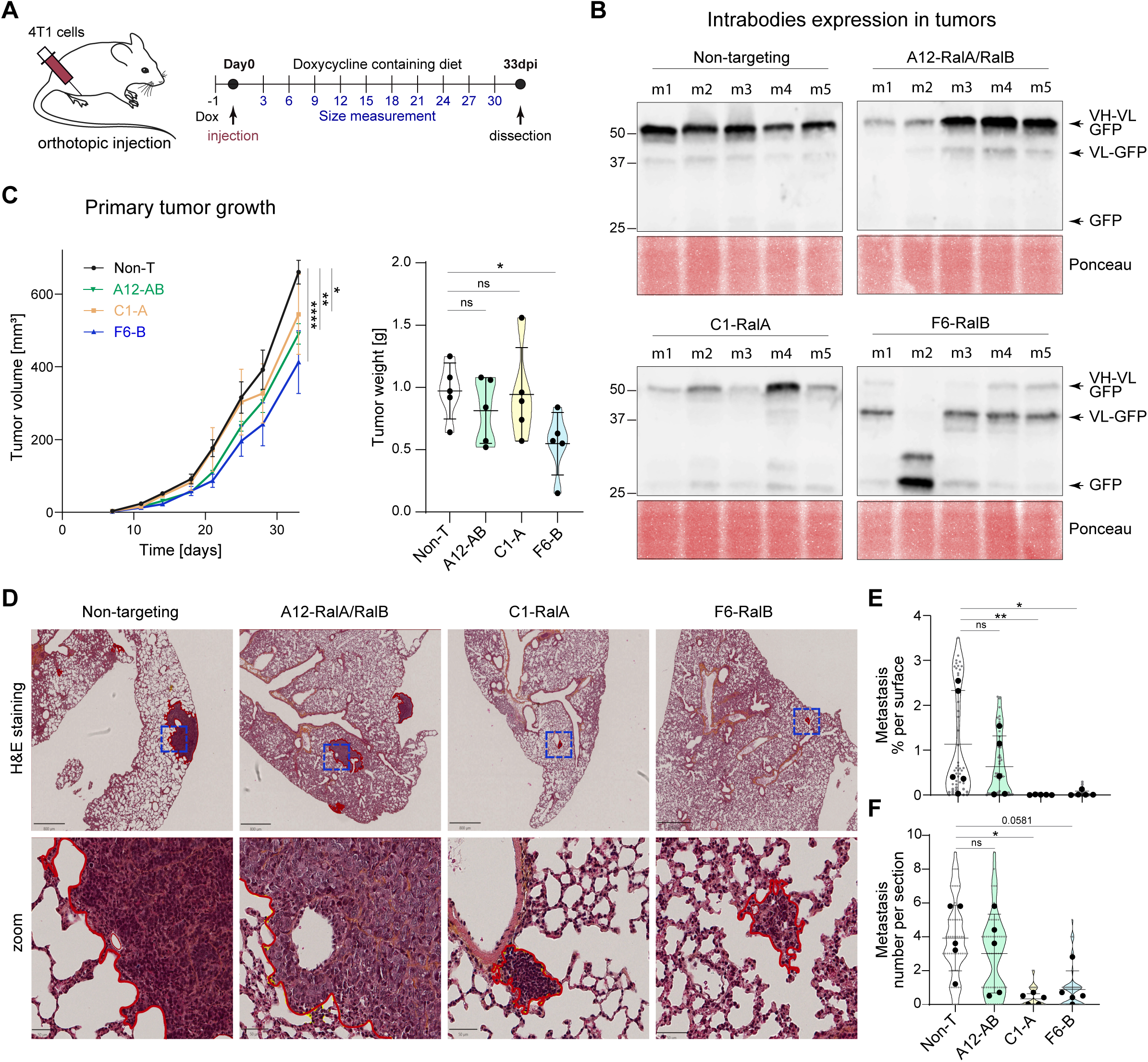
Impact of intrabody expression on primary tumor growth and metastasis. **A**) 4T1 cells stably expressing non-targeting (Non-T), pan-specific (A12-AB), RalA-specific (C1-A) and RalB-specific (F6-B) intrabodies were injected orthotopically in nude mice. Cells were treated with doxycycline (1 µg/ml) 24 h before injection and mice were kept on doxycycline diet. Tumor volume was measured every 3 days up to 33 days post injection. **B**) Western blot analysis of GFP expression in the primary tumors (Day 33). Arrows indicate, from top to bottom, full-length scFv-GFP, VL-GFP (cleavage in the scFv linker), and free GFP. Bottom (red membrane stain): total protein loading. **C**) Left: Graph showing the primary tumor growth over time until day 33 (n = 5 mice per condition). Tumor volume at Day 33 of 4T1 cells expressing Non-T, A12-AB, C1-A and F6-B clone (Mean ± SD): 660.4 ± 73.1, 490.5 ± 62.5, 544.4 ± 246.4, 412.9 ± 192.8. Two-way ANOVA with two-stage linear step-up procedure of Benjamini, Krieger and Yekutieli for multiple comparisons. Right: Primary tumor weight at day 33 for 4T1 cells expressing Non-T, A12-AB, C1-A and F6-B clone (Mean ± SD): 0.9720 ± 0.2247, 0.8140 ± 0.2636, 0.9440 ± 0.3755, 0.5480 ± 0.2504. Kruskal-Wallis with Dunn’s Multiple Comparison Test. **D**) Analysis of lung metastasis in mice, orthotopic model. Representative images of lung sections at day 33, Hematoxylin / Eosin / Safran staining. Scale bar = 800 µm, zoom = 50 µm, red line= metastasis outline. **E**) Metastasis surface in lungs (n = 10 sections / mouse). Small gray dots represent % of metastasis per lung surface for each section. Black dots represent average value per mouse (Non-T, A12-AB, C1-A, F6-B clone; Mean ± SD): 1.137 ± 1.203, 0.6336 ± 0.6878, 0.0060 ± 0.0060, 0.0318 ± 0.0556. Statistical analysis, Kruskal-Wallis with Dunn’s Multiple Comparison Test. **F**) Number of metastatic foci per section (small gray dots) and average number of metastatic foci per mouse (black dots). Number of foci (Non-T, A12-AB, C1-A, F6-B clone; Mean ± SD): 3.920 ± 1.942, 3.000 ± 2.329, 0.3208 ± 0.3120, 0.8800 ± 1.103. Statistical analysis: Kruskal-Wallis with Dunn’s Multiple Comparison Test. * p□<□0.05; ** p□<□0.01; *** p□<□0.001; **** p□<□0.0001.

When analyzing the impact of anti-Ral intrabodies on lung metastasis (Fig. 5D, 5E & 5F), we observed that both anti-RalA C1-A and anti-RalB F6-B intrabodies strongly decreased both the number of metastatic foci and their total surface. Notably, a complete absence of detectable metastases was observed in 2 out of 5 mice treated with C1-A and in 1 out of 5 mice treated with F6-B, underscoring the potent anti-metastatic effect of these intrabodies. By contrast, metastases developed similarly in tumors expressing the pan-Ral intrabody A12-AB as in tumors expressing the control non-T intrabody. Altogether, we have generated intrabodies targeting specific Ral isoforms which are able to strongly inhibit metastasis *in vivo*.

## Discussion

In this study, we characterized two intrabodies specifically recognizing the RalA small GTPase. The intrabody approach was chosen due to the lack of available pharmacological tools for specifically targeting RalA paralog. Both intrabodies, expressed as scFv, demonstrated high in vitro specificity by ELISA (Fig. 1B). Despite their relatively large size (30 kDa) compared to RalA (23 kDa), the two intrabodies recognize close but distinct epitopes, as evidenced by the partial binding competition between the two clones (Fig. 1D & Supplementary Fig. S3A). They also did not compete with a third pan-RalAB clone (A12-AB), showing that at least 3 independent epitopes can be targeted on RalA.

When expressed as GFP fusions in murine 4T1 cells, C1-A and A12-AB intrabodies were localized to the cytoplasm and co-localized with their RalA target, as shown by immunofluorescence studies (Fig. 2F & 2G), but this was less visible with G5-A because of the instability of the link between the scFv and the GFP tracer. We further demonstrated their intracellular activity by characterizing their effect on mitochondrial morphology (Fig. 4). All RalA-targeting intrabodies induced an increase in mitochondrial length, a phenotype associated with impaired mitochondrial fission and previously observed in RalA-depleted cells using shRNA (Kashatus *et al*, 2011; Ghoroghi *et al*, 2021). This effect was not observed after transduction with the RalB-targeting F6-B intrabody and the non-T control, which maintained normal mitochondrial morphology. Interestingly, A12-AB pan-Ral intrabody, recognizing an epitope different from the two RalA-specific clones, also induced this mitochondrial effect. This demonstrates that the mitochondrial phenotype can be achieved by targeting various sites on the RalA protein.

The two RalA-specific intrabodies exhibited distinct effects on the endo-lysosomal system (Fig. 3). Specifically, the number of lysosomes was significantly decreased by both intrabodies, but lysosome size was only affected by C1-A. Of note, despite recognizing RalA, the pan-Ral intrabody A12-AB did not modify the endo-lysosomal system. Furthermore, 4T1 cells expressing the C1-A intrabody showed a marked decrease in the number of extracellular vesicles (EVs) in the cell supernatant. This specific reduction in EV number was not observed with other RalA-targeting clones (G5-1 and A12-AB) or with the RalB-specific clone F6-B. The distinct effects on the endo-lysosomal system highlight the epitope-specific action of intrabodies, contrary to the shRNA-based approach that silence all phenotypes associated with RalA (Ghoroghi *et al*, 2021).

Most notably, the C1-A intrabody strongly inhibited metastasis formation and growth in an orthotopic breast cancer model (Fig. 5D, 5E & 5F). Indeed, 2 out of 5 (40%) mice showed no metastasis at the end of the experiment (Fig. 5F), and the remaining 3 mice (60%) exhibited a four-fold reduction in metastasis number and a ten-fold reduction in metastatic lesions compared to a non-targeting intrabody. This pronounced anti-metastatic effect contrasts sharply with the weak impact on primary tumor growth (Fig. 5C). In addition, while shRNA depletion in 4T1 mice resulted in an increase in primary tumor growth and a decrease of metastases (Ghoroghi *et al*, 2021), C1-A intrabody inhibited both phenotypes and resulted in a much deeper reduction of metastasis formation and size. This demonstrated that pharmacological-like action via specific binding rather than complete target depletion can be much more efficient for metastasis reduction. In contrast, the RalB-specific clone F6-B affected both primary tumor growth and metastasis number and size, albeit to a lesser extent than C1-A for metastasis, and closely matched shRNA knock down.

While the precise mode of action of RalA-targeting intrabodies remains to be fully elucidated, our results suggest that they do not function by depleting their target, as the observed phenotypes differ across the three RalA-recognizing clones targeting three different epitopes on the protein, and because they are only partially comparable to previously described shRNA-based depletions. Furthermore, the intrabodies do not appear to induce RalA mislocalization, as we did not detect any change in RalA localization or expression (Fig. 2F). The decrease in metastasis formation could be linked to the strong and specific inhibition of EV secretion in C1-A expressing cells, a phenomenon also observed in RalA-depleted cells using shRNA (Ghoroghi *et al*, 2021). Extracellular vesicles are known to travel from primary tumor sites via blood vessels and the lymphatic system, remodeling the metastatic niche and promoting cell implantation and metastatic growth (Becker *et al*, 2016). Despite an unknown mechanism, the dependency of cell phenotypes on the targeted epitope demonstrates that specific inhibition of particular RalA activities can be achieved through pharmacological modulation using intrabodies, leading, in the case of C1-A, to an almost complete blockade of metastasis formation and growth, independently of primary tumor growth rate.

Over the past years, intrabodies have emerged as promising new therapeutic tools, particularly in cancer (Lin *et al*, 2020). While this tool is primarily restricted to *in vitro* studies and preclinical experiments, some data support their clinical use. A clinical trial using adenoviral delivery of an intrabody directed against Her2 in human ovarian cancer demonstrated the safety and manageability of such therapies (Alvarez *et al*, 2000). Additionally, several reports have used viral methods, in particular adenovirus, adenovirus-related viruses and lentivirus, to deliver genes encoding intrabodies in vivo (Lin *et al*, 2019). Finally, antibodies can be modified to enter the cell, escape lysosome, and target proteins in the cytoplasm, the nucleus or any other cell compartment (Shin *et al*, 2017). As an alternative to delivery, a highly promising approach is to leverage data obtained with intrabodies to design or select small chemical drugs that can be produced at lower cost and delivered orally (Stocks, 2005; Lawson, 2012). Two main strategies have been developed: the use of intrabody chemical mimics identified through competition experiments, as illustrated with Syk tyrosine kinase (Mazuc *et al*, 2008; Villoutreix *et al*, 2011) or RAS oncogene (Quevedo *et al*, 2018), and the use of antibody to constrain a protein in an inactive state, thereby exposing new binding pockets for drugs (Mullard, 2022), as demonstrated for KRAS (Davies *et al*, 2022) or GPCR (Pardon *et al*, 2018). Future experiments will be essential to clearly identify the precise mechanism of action of the RalA-targeting intrabodies described here, and if this could help in the design of pharmacological drugs targeting RalA and inhibiting metastasis formation.

## Materials and methods

### scFv selection by phage display

The human synthetic HuscI2 library in scFv format is based on the sequence of the optimized hyper-stable 13R4 scFv (Martineau *et al*, 1998; Martineau & Betton, 1999) and is displayed on filamentous phages using the pHEN1 vector. Diversity was introduced at positions [Aho numbering (Honegger & Plückthun, 2001)] L38-40, L58, L69, L109, L111, L112, H39-40, H57, H59, H67, H69, and with eight 2-9 residue loops inserted between positions H110-134 of VH-CDR3 (Robin *et al*, 2014). Positions were diversified with a mix of five amino acids (Y, S, D, N, G) by gene synthesis (Twist Bioscience) independently in VH and VL sequences. To limit liabilities in the resulting antibodies, sites of N-glycosylation (NxS/T), NG (N deamidation) and DG (D isomerization) sequences (Lu *et al*, 2018) were not included in the library and replaced, depending on the context, by NxA or QxS/T, NA or QG, and DA or EG, respectively. The final library was assembled by Golden-Gate cloning and transformed in TG1 bacteria by electroporation. According to the number of colonies, the clonal diversity is 3 × 10^10^. Frequencies of mutations and correctness of the genes (> 95%) were confirmed by NGS sequencing on Illumina platform.

The library was screened to select anti-RalA and anti-RalB specific scFvs by phage display according to standard protocols (Philibert *et al*, 2007). Briefly, 4 rounds of panning were performed with or without an initial depletion on Ral paralog. Individual secreted scFv were screened by ELISA. Antigens, for depletion and selection, were captured at 10 µg/mL on 96-well MaxiSorp immunoplates (Nunc) coated with streptavidin and incubated with Tris-NTA-biotin (biotechrabbit GmbH # BR1001201).

### Production and purification of recombinant proteins

Human recombinant 6xHis tagged RalA and RalB were produced in BL21(DE3)-pLysS as previously described (Diskin *et al*, 2004). Bacteria were grown in 2xTY medium with 100 µg/mL ampicillin and when culture reached OD_600_ = 0.5, 6xHis-fusion protein expressions were induced using 1 mM isopropyl β-D-1-thiogalactopyranoside (IPTG) at 30°C for at least 4h. Bacteria were harvested, then frozen at -20°C overnight. Thawed pellets were resuspended in a lysis buffer (10 mM HEPES pH8, 0.5 mM EDTA, 30 mM NaCl, 0.65% NP-40, Benzonase 5 U/mL) and supernatants were clarified by successive centrifugations.

ScFvs tagged with a c-Myc /Flag and a 6xHis tag at their C-terminus were cloned in a pAB1 vector and expressed in the periplasm of HB2151 strain. Cells were grown in 2xTY medium with 100 µg/mL ampicillin at 37°C until cultures reached OD_600_ = 0.5-0.7, 1 mM. Subsequently IPTG was added and growth was continued at 30°C for at least 4 h. Then bacteria were harvested by centrifugation. The cells pellets were resuspended in a hypertonic buffer (Tris-HCl 30mM pH 8, EDTA 1 mM, saccharose 20%) in a volume of 1/20 of the culture volume, incubated 30 min on ice and then centrifuged 20 min at 6,000 x g. The second pellets were resuspended in the same volume of a hypotonic buffer (MgCl2 1 mM), incubated 30 min on ice and then centrifugated 20 min at 6,000 x g. The supernatants were clarified again by centrifugation. After bacteria lysis, 6xHis tagged proteins were purified by affinity chromatography on HisPur™ Cobalt resin (Thermo Scientific # 89965).

### Competitive ELISA assay

Antigens in PBS were coated at 2 and 4 µg/mL for RalA and RalB, respectively, in 96-well MaxiSorp immunoplates (Nunc) overnight at 4 °C. Wells were blocked 1 hour at RT with 5 % milk in PBS with Tween20 0.1 %. C-Myc-tagged scFvs were added at a saturating concentration of 30 µg/mL then a detection was done with a Flag-tagged scFvs at a concentration giving a signal linearly proportional to the concentration (2 µg/mL) during 2 hours. Signal was detected with a peroxidase-conjugated anti-Flag antibody (Sigma Aldrich, # A8592).

### Lentivirus production and infection

Stable cell lines expressing the different scFv were obtained by lentiviral gene transduction. To this aim, scFv genes were cloned in PCLX plasmid. Lentiviral particles were produced in 293T by transient co-transfection of gag/pol, env and the viral PCLX constructs. More precisely, cells were seeded and infect with PCLX-scFv vector on jetPEI transfection reagent (Polyplus TransfectionTM) were diluted in 150 mM NaCl. After 5 h, the medium was replaced with 10 ml of fresh medium. The supernatant containing the virus was collected 72 h later, precipitated with PEG-it™ Virus Precipitation solution and conserved at -80°C. To obtain stable cell lines expressing scFvs, lentiviruses were used to transduce the 4T1 cells seeded in a 6 well plate, 400,000 cells/well, in the presence of 5µg/mL of polybrene (Sigma Aldrich, reference H9268). 48h later, the transduced cells were selected with blasticidin at 5 µg/mL (Sigma Aldrich, reference 15205-100MG).

### Cell culture

Cells were cultured at 37 °C with 5% CO2 in DMEM (HEK 293-T) or RPMI (4T1 cell) medium containing 10% FBS. Blasticidine at 5µM was added in 4T1 culture medium of cells stably expressing scFv-GFP to maintain the selection pressure. The Doxycycline was added at 0.5 μg/ml to the media at different timing to induce expression of the TetOn promoter and scFv-GFP production.

### Plasmid transfections

Cells at 50–70% confluency were transfected with 1 μg of pLenti CMV:tdtomato-RalA, and pLenti CMV:tdtomato-RalB plasmid using JetPRIME (PolyPlus, Illkirch, France) according to the manufacturer’s instructions.

### Cytometry

Cells were seeded in 6-well plate at 50,000 cells by well at Day 0 and were grown for 72 h. To induce intrabodies expression, cells were treated with Dox for 48, 24 or 6 h. Cells were then washed, centrifuged and fixed in 2% PFA. The GFP fluorescence was measured by flow cytometry with the Cytoflex LX cytometer (Beckman coulter). Flow cytometry data were quantified using the FlowJo-V10 software.

### Western blot

Cells were lysed at 4 °C in RIPA lysis buffer with 1% protease and phosphatase inhibitor (ThermoFisher 78430). Laemmli buffer (4% SDS, 20% glycerol, 1% 2-β mercaptoethanol, 0.004 % bromophenol blue, 0.125 mol/L Tris HCL) with benzonase (Millipore # 70746-3) was then added to the whole cell lysate and boiled at 95 °C for 5 minutes. Cell lysate was loaded on agarose gel 12% acrylamide and then transferred on 0.45 μm nitrocellulose membranes. Blocking was done with 5% BSA in TBS-tween for 2 h at RT. Membranes were incubated with primary antibodies overnight at 4 °C, washed, and then incubated with secondary antibodies for 2h at RT. Detection was performed alternatively with enhanced chemiluminescence method (Western Lightning Plus-ECL, PerkinElmer) when using HRP-conjugated secondary antibodies or with the LICOR Odyssey Infrared Imaging System (LI-COR Biosciences) when using fluorescent-conjugated secondary antibodies (IRDye® 800CW Goat anti-Rabbit IgG, LICOR #926-32211, 1/10,000). GFP: Invitrogen #A11122 1/1000 ; His-tag : Sigma Aldrich #A7058 1/2000; c-myc-tag: 9E10 clone, sc40-HRP Santa Cruz Biotechnology 1/200; Alix (BD Biosciences #611621 1/1000), CD9 (BD Biosciences #553758 1/1000) and TSG101 (GeneTex, #GTX70255; 1/1000).

### Fluorescence microscopy

Cells were plated in 4-chamber glass bottom dishes (35 mm, #1.5, Cellvis). To assess Mitochondrial network and lysosome number, cells were stained with respectively 200 nM of Mitotracker deep red (Molecular Probes #M22426) or 50 nM Green Lysotracker (Molecular Probes #L7526) for 20 minutes before being imaged using Olympus Spinning Disk (60X objective, N.A. 1.2), 37°C and 5% CO2. Image analysis and processing were performed using Fiji (v2.0.0) for mitochondrial aspect ratio quantification (Schindelin et al., 2012). Mitochondria were segmented from background using thresholding and binary processing tools. Individual organelles were analyzed using the “Analyze Particles” function. Aspect ratio was calculated as the ratio of the major to minor axis of each mitochondrion to assess changes in morphology. To assess lysosome parameters, cells imaged after LysoTracker staining, were processed through a custom pipeline to identify and measure lysosomal objects based on their size on Cell Profiler (4.2.1) software. In short, lysosome objects were identified by thresholding, their size corresponds to the number of pixels in each object per cell.

### Electron microscopy

Cells were fixed with 2.5 % glutaraldehyde / 2.0 % paraformaldehyde (PFA) (Electron Microscopy Sciences) in 0.1 M Cacodylate buffer at room temperature for 2 hours, then rinsed in 0.1 M Cacodylate buffer (Electron Microscopy Sciences) and post-fixed with 1% OsO4 (Electron Microscopy Sciences) and 0.8% K_3_Fe(CN)_6_ (Sigma-Aldrich) for 1 hour at 4°C. Then, samples were rinsed in 0.1 M Cacodylate buffer followed by a water rinse and stained with 1 % uranyl acetate, overnight at 4 °C. The samples were stepwise dehydrated in Ethanol (50 %, 70 % 2 × 10 min, 95 % 2 × 15 min and 100 % 3 × 15 min), infiltrated in a graded series of Epon (Ethanol 100 % / Epon 3/1, 1/1, 1 h) and kept in Ethanol 100 % / Epon 1/3 overnight. The following day, samples were placed in pure Epon and polymerized at 60°C. One hundred nm thin sections were collected in 200 copper mesh grids and imaged with a Philips CM12 transmission electron microscope operated at 80 kV and equipped with an Orius 1000 CCD camera (Gatan).

### EV isolation, quantification, and characterization

Cells were cultured in EV-depleted medium (obtained by overnight ultracentrifugation at 100,000 g, using a Beckman, XL-70 centrifuge with a 70Ti rotor) for 24 h before supernatant collection. EVs were isolated by successive centrifugation at 4 °C: 15 min at 300 g, 10 min at 2000 g, 30 min at 10,000 g and 70 min at 100,000 g (using a Beckman XL-70 centrifuge with a SW28 rotor). EVs pellets were washed in PBS, centrifuged again at 100,000 g for 70 min, resuspended in PBS quantified by NTA using a ZetaView (Particle Metrix, Meerbusch, Germany). EVs were stored at -20 °C and characterized by WB.

### Animal experiments

All animals were housed and handled according to the guidelines of INSERM and the ethical committee of Alsace, France (CREMEAS) (Directive 2010/63/EU on the protection of animals used for scientific purposes). Animal facility agreement number: #C67-482-33. Experimental license for mice: Apafis #4707 et #38090). Eight-week-old female nude mice (Crl:NU(Ncr)-Foxn1^nu^, Charles River) were grafted Orthotopically in the left fourth mammary gland with 250,000 4T1 mammary tumor cells expressing each intrabody. Mice were fed with Doxycycline-containing food (1 g/kg; Safe, #E8200) during the whole procedure to maintain scFvs expression. Tumor volume was assessed every 3 days by caliper measurements using the formula (width^2^ × length)/2 [mm^3^]. At the endpoint of the experiment, tumors were harvested, weighted and frozen at -80 °C. Mouse lungs were incubated overnight in 4 % PFA, dehydrated in 100 % ethanol for 24 h, embedded in paraffin, cut in 7μm thick sections, dewaxed and rehydrated with 100 % Toluene (2 washes of 15 min) then incubated in 100 %-70 % alcohol solutions (10 min each) followed by final staining with hematoxylin (Surgipath) for 5 min and washing with tap water. Sections were further processed with differentiation solution (1 % HCl in absolute ethanol, for 7 s), followed by washing under tap water for 10 min. Sections were then incubated in eosin (Harris) for 10 s, rinsed and dehydrated in 70 % - 100 % alcohol baths with rapid dips in each bath before a final wash in toluene for 15 min and embedded in Eukitt solution (Sigma). Ten 3 µm sections with a 15 µm gap between the different sections taken from each lung were analyzed for each mouse. Sections were imaged using a Nanozoomer 2 Hamamatsu. Metastasis quantification was performed using QuPath (0.5.0) open-source software. Regions of interest (ROIs) were annotated semi-automatically, and metastatic foci were quantified based on defined morphological and staining criteria. Data were then exported for statistical analysis on GraphPad Prism.

## Acknowledgements

This project was conducted within the NANOTUMOR Consortium, a program from ITMO Cancer of AVIESAN (Alliance Nationale pour les Sciences de la Vie et de la Santé, National Alliance for Life Sciences & Health; ASC19052FSA) co-headed by JGG within the framework of the Cancer Plan (France). We are particularly grateful to Florent Colin who coordinated the Nanotumor consortium. CS and KJR were funded by Nanotumor, KJR was also supported by FRM (post-doctoral fellowship SPF202004011876). Work and people in the Tumor Biomechanics Lab are mostly supported by the INCa (Institut National Du Cancer, French National Cancer Institute), charities (La Ligue contre le Cancer, ARC (Association pour la Recherche contre le Cancer), FRM (Fondation pour la Recherche Médicale), the National Plan Cancer initiative, the Region Grand Est, INSERM and the University of Strasbourg and from local donators (TDLR, Club Féminin Lampertheim). This work has been directly funded by the support of the Ligue Contre le Cancer (labelisation) and the association Ruban Rose, with additional support from INCa. LB has been funded by FRM (Fondation pour la Recherche Médicale) and CL by INCa. We thank G. Khelifi and C. Hergott for animal care. Work in Functional screening and targeting in cancer team has been partially funded with support from the French National Research Agency under the program “Investissements d’avenir” Grant Agreement LabEx MAbImprove: ANR-10-LABX-53. The imaging was supported by PIC-STRA (CRBS, Strasbourg: P. Kessler, fluorescent imaging) and PIV (INCI, Strasbourg: C. Royer, V. Demais, electron microscopy) imaging platforms, members of the national infrastructure France-BioImaging supported by the French National Research Agency (ANR-10-INBS-04). We thank all members of the Tumor Biomechanics Lab for their constant helpful discussions throughout this investigation.

## References

Abba Moussa D, Vazquez M, Chable-Bessia C, Roux-Portalez V, Tamagnini E, Pedotti M, Simonelli L, Ngo G, Souchard M, Lyonnais S, Chentouf M, Gros N, Marsile-Medun S, Dinter H, Pugnière M, Martineau P, Varani L, Juan M, Calderon H, Naranjo-Gomez M, Pelegrin M (2025) Discovery of a pan anti-SARS-CoV-2 monoclonal antibody with highly efficient infected cell killing capacity for novel immunotherapeutic approaches. Emerg Microbes Infect 14: 2432345, doi:10.1080/22221751.2024.2432345.

Alvarez RD, Barnes MN, Gomez-Navarro J, Wang M, Strong TV, Arafat W, Arani RB, Johnson MR, Roberts BL, Siegal GP, Curiel DT (2000) A cancer gene therapy approach utilizing an anti-erbB-2 single-chain antibody-encoding adenovirus (AD21): a phase I trial. Clin Cancer Res Off J Am Assoc Cancer Res 6: 3081–3087.

Apken LH, Oeckinghaus A (2021) Chapter Two - The RAL signaling network: Cancer and beyond. In International Review of Cell and Molecular Biology, T.S. Postler, and L. Galluzzi, eds (Academic Press), pp. 21–105.

Becker A, Thakur BK, Weiss JM, Kim HS, Peinado H, Lyden D (2016) Extracellular vesicles in cancer: cell-to-cell mediators of metastasis. Cancer Cell 30: 836–848, doi:10.1016/j.ccell.2016.10.009.

Cattaneo A, Chirichella M (2019) Targeting the Post-translational Proteome with Intrabodies. Trends Biotechnol 37: 578–591, doi:10.1016/j.tibtech.2018.11.009.

Davies CW, Oh AJ, Mroue R, Steffek M, Bruning JM, Xiao Y, Feng S, Jayakar S, Chan E, Arumugam V, Uribe SC, Drummond J, Frommlet A, Lu C, Franke Y, Merchant M, Koeppen H, Quinn JG, Malhotra S, Do S, Gazzard L, Purkey HE, Rudolph J, Mulvihill MM, Koerber JT, Wang W, Evangelista M (2022) Conformation-locking antibodies for the discovery and characterization of KRAS inhibitors. Nat Biotechnol 40: 769–778, doi:10.1038/s41587-021-01126-9.

De Groof TWM, Bergkamp ND, Heukers R, Giap T, Bebelman MP, Goeij-de Haas R, Piersma SR, Jimenez CR, Garcia KC, Ploegh HL, Siderius M, Smit MJ (2021) Selective targeting of ligand-dependent and -independent signaling by GPCR conformation-specific anti-US28 intrabodies. Nat Commun 12: 4357, doi:10.1038/s41467-021-24574-y.

Diskin R, Askari N, Capone R, Engelberg D, Livnah O (2004) Active mutants of the human p38alpha mitogen-activated protein kinase. J Biol Chem 279: 47040–47049, doi:10.1074/jbc.M404595200.

Douglass J, Hsiue EH-C, Mog BJ, Hwang MS, DiNapoli SR, Pearlman AH, Miller MS, Wright KM, Azurmendi PA, Wang Q, Paul S, Schaefer A, Skora AD, Molin MD, Konig MF, Liu Q, Watson E, Li Y, Murphy MB, Pardoll DM, Bettegowda C, Papadopoulos N, Gabelli SB, Kinzler KW, Vogelstein B, Zhou S (2021) Bispecific antibodies targeting mutant RAS neoantigens. Sci Immunol 6: eabd5515, doi:10.1126/sciimmunol.abd5515.

Falsetti SC, Wang D, Peng H, Carrico D, Cox AD, Der CJ, Hamilton AD, Sebti SM (2007) Geranylgeranyltransferase I inhibitors target RalB to inhibit anchorage-dependent growth and induce apoptosis and RalA to inhibit anchorage-independent growth. Mol Cell Biol 27: 8003–8014, doi:10.1128/MCB.00057-07.

Ghoroghi S, Mary B, Larnicol A, Asokan N, Klein A, Osmani N, Busnelli I, Delalande F, Paul N, Halary S, Gros F, Fouillen L, Haeberle A-M, Royer C, Spiegelhalter C, André-Grégoire G, Mittelheisser V, Detappe A, Murphy K, Timpson P, Carapito R, Blot-Chabaud M, Gavard J, Carapito C, Vitale N, Lefebvre O, Goetz JG, Hyenne V (2021) Ral GTPases promote breast cancer metastasis by controlling biogenesis and organ targeting of exosomes. eLife 10: e61539, doi:10.7554/eLife.61539.

Guglielmi L, Denis V, Vezzio-Vié N, Bec N, Dariavach P, Larroque C, Martineau P (2011) Selection for intrabody solubility in mammalian cells using GFP fusions. Protein Eng Des Sel PEDS 24: 873–881, doi:10.1093/protein/gzr049.

Guin S, Ru Y, Wynes MW, Mishra R, Lu X, Owens C, Barόn AE, Vasu VT, Hirsch FR, Kern JA, Theodorescu D (2013) Contributions of KRAS and RAL in Non-Small Cell Lung Cancer growth and progression. J Thorac Oncol Off Publ Int Assoc Study Lung Cancer 8: 1492–1501, doi:10.1097/JTO.0000000000000007.

Han D, Spehar JM, Richardson DS, Leelananda S, Chakravarthy P, Grecco S, Reardon J, Stover DG, Bennett C, Sizemore GM, Li Z, Lindert S, Sizemore ST (2024) The RAL Small G Proteins Are Clinically Relevant Targets in Triple Negative Breast Cancer. Cancers 16: 3043, doi:10.3390/cancers16173043.

Honegger A, Plückthun A (2001) Yet Another Numbering Scheme for Immunoglobulin Variable Domains: An Automatic Modeling and Analysis Tool. J Mol Biol 309: 657–670, doi:10.1006/jmbi.2001.4662.

Hsiue EH-C, Wright KM, Douglass J, Hwang MS, Mog BJ, Pearlman AH, Paul S, DiNapoli SR, Konig MF, Wang Q, Schaefer A, Miller MS, Skora AD, Azurmendi PA, Murphy MB, Liu Q, Watson E, Li Y, Pardoll DM, Bettegowda C, Papadopoulos N, Kinzler KW, Vogelstein B, Gabelli SB, Zhou S (2021) Targeting a neoantigen derived from a common TP53 mutation. Science 371: eabc8697, doi:10.1126/science.abc8697.

Hurd CA, Brear P, Revell J, Ross S, Mott HR, Owen D (2021) Affinity maturation of the RLIP76 Ral binding domain to inform the design of stapled peptides targeting the Ral GTPases. J Biol Chem 296: doi:10.1074/jbc.RA120.015735.

Hyenne V, Apaydin A, Rodriguez D, Spiegelhalter C, Hoff-Yoessle S, Diem M, Tak S, Lefebvre O, Schwab Y, Goetz JG, Labouesse M (2015) RAL-1 controls multivesicular body biogenesis and exosome secretion. J Cell Biol 211: 27–37, doi:10.1083/jcb.201504136.

Kashatus DF, Lim K-H, Brady DC, Pershing NLK, Cox AD, Counter CM (2011) RALA and RALBP1 regulate mitochondrial fission at mitosis. Nat Cell Biol 13: 1108–1115, doi:10.1038/ncb2310.

Lawson ADG (2012) Antibody-enabled small-molecule drug discovery. Nat Rev Drug Discov 11: 519–525, doi:10.1038/nrd3756.

Lin X-R, Zhou X-L, Feng Q, Pan X-Y, Song S-L, Fang H, Lei J, Yang J-L (2019) CIK cell-based delivery of recombinant adenovirus KGHV500 carrying the anti-p21Ras scFv gene enhances the anti-tumor effect and safety in lung cancer. J Cancer Res Clin Oncol 145: 1123–1132, doi:10.1007/s00432-019-02857-8.

Lin Y, Chen Z, Hu C, Chen Z-S, Zhang L (2020) Recent progress in antitumor functions of the intracellular antibodies. Drug Discov Today 25: 1109–1120, doi:10.1016/j.drudis.2020.02.009.

Lu X, Nobrega RP, Lynaugh H, Jain T, Barlow K, Boland T, Sivasubramanian A, Vásquez M, Xu Y (2018) Deamidation and isomerization liability analysis of 131 clinical-stage antibodies. mAbs 11: 45–57, doi:10.1080/19420862.2018.1548233.

Martineau P, Betton JM (1999) In vitro folding and thermodynamic stability of an antibody fragment selected in vivo for high expression levels in Escherichia coli cytoplasm. J Mol Biol 292: 921–929.

Martineau P, Jones P, Winter G (1998) Expression of an antibody fragment at high levels in the bacterial cytoplasm. J Mol Biol 280: 117–127.

Mazuc E, Villoutreix BO, Malbec O, Roumier T, Fleury S, Leonetti J-P, Dombrowicz D, Daëron M, Martineau P, Dariavach P (2008) A novel druglike spleen tyrosine kinase binder prevents anaphylactic shock when administered orally. J Allergy Clin Immunol 122: 188–194, 194.e1-3, doi:10.1016/j.jaci.2008.04.026.

Mullard A (2022) Antibody clamps pry open small-molecule drug discovery opportunities. Nat Rev Drug Discov 21: 247–248, doi:10.1038/d41573-022-00054-w.

Pardon E, Betti C, Laeremans T, Chevillard F, Guillemyn K, Kolb P, Ballet S, Steyaert J (2018) Nanobody-Enabled Reverse Pharmacology on G-Protein-Coupled Receptors. Angew Chem Int Ed Engl 57: 5292–5295, doi:10.1002/anie.201712581.

Peschard P, McCarthy A, Leblanc-Dominguez V, Yeo M, Guichard S, Stamp G, Marshall CJ (2012) Genetic Deletion of RALA and RALB Small GTPases Reveals Redundant Functions in Development and Tumorigenesis. Curr Biol 22: 2063–2068, doi:10.1016/j.cub.2012.09.013.

Philibert P, Stoessel A, Wang W, Sibler A-P, Bec N, Larroque C, Saven JG, Courtête J, Weiss E, Martineau P (2007) A focused antibody library for selecting scFvs expressed at high levels in the cytoplasm. BMC Biotechnol 7: 81, doi:10.1186/1472-6750-7-81.

Quevedo CE, Cruz-Migoni A, Bery N, Miller A, Tanaka T, Petch D, Bataille CJR, Lee LYW, Fallon PS, Tulmin H, Ehebauer MT, Fernandez-Fuentes N, Russell AJ, Carr SB, Phillips SEV, Rabbitts TH (2018) Small molecule inhibitors of RAS-effector protein interactions derived using an intracellular antibody fragment. Nat Commun 9: 3169, doi:10.1038/s41467-018-05707-2.

Richardson DS, Spehar JM, Han DT, Chakravarthy PA, Sizemore ST (2022) The RAL Enigma: Distinct Roles of RALA and RALB in Cancer. Cells 11: 1645, doi:10.3390/cells11101645.

Robin G, Sato Y, Desplancq D, Rochel N, Weiss E, Martineau P (2014) Restricted diversity of antigen binding residues of antibodies revealed by computational alanine scanning of 227 antibody-antigen complexes. J Mol Biol 426: 3729–3743, doi:10.1016/j.jmb.2014.08.013.

Shin S-M, Choi D-K, Jung K, Bae J, Kim J-S, Park S-W, Song K-H, Kim Y-S (2017) Antibody targeting intracellular oncogenic Ras mutants exerts anti-tumour effects after systemic administration. Nat Commun 8: 15090, doi:10.1038/ncomms15090.

Sibler A-P, Nordhammer A, Masson M, Martineau P, Trave G, Weiss E (2003) Nucleocytoplasmic shuttling of antigen in mammalian cells conferred by a soluble versus insoluble single-chain antibody fragment equipped with import/export signals. Exp Cell Res 286: 276–287.

Stocks M (2005) Intrabodies as drug discovery tools and therapeutics. Curr Opin Chem Biol 9: 359–365.

Thies KA, Cole MW, Schafer RE, Spehar JM, Richardson DS, Steck SA, Das M, Lian AW, Ray A, Shakya R, Knoblaugh SE, Timmers CD, Ostrowski MC, Chakravarti A, Sizemore GM, Sizemore ST (2021) The small G-protein RalA promotes progression and metastasis of triple-negative breast cancer. Breast Cancer Res BCR 23: 65, doi:10.1186/s13058-021-01438-3.

Thomas JC, Cooper JM, Clayton NS, Wang C, White MA, Abell C, Owen D, Mott HR (2016) Inhibition of Ral GTPases Using a Stapled Peptide Approach. J Biol Chem 291: 18310–18325, doi:10.1074/jbc.M116.720243.

Villoutreix BO, Laconde G, Lagorce D, Martineau P, Miteva MA, Dariavach P (2011) Tyrosine kinase syk non-enzymatic inhibitors and potential anti-allergic drug-like compounds discovered by virtual and in vitro screening. PloS One 6: e21117, doi:10.1371/journal.pone.0021117.

Walsh TG, Wersäll A, Poole AW (2019) Characterisation of the Ral GTPase inhibitor RBC8 in human and mouse platelets. Cell Signal 59: 34–40, doi:10.1016/j.cellsig.2019.03.015.

Yan C, Liu D, Li L, Wempe MF, Guin S, Khanna M, Meier J, Hoffman B, Owens C, Wysoczynski CL, Nitz MD, Knabe EW, Brautigan DL, Paschal BM, Schwartz MA, Jones D, Ross D, Meroueh SO, Theodorescu D (2014) Discovery and characterization of small molecules that target the Ral GTPase. Nature 515: 443–447, doi:10.1038/nature13713.

Yan C, Theodorescu D (2018) RAL GTPases: Biology and Potential as Therapeutic Targets in Cancer. Pharmacol Rev 70: 1–11, doi:10.1124/pr.117.014415.

