## Supplementary Figures for "Selective targeting of RalA with intrabodies impairs Triple Negative Breast Cancer metastasis"

```
                10        20        30        40        50        60        70
        ----:----|----:----|----:----|----:----|----:----|----:----|----:----|
fwk     MAEVQLVESGGGLVKPGGSLRLSCAASGFTFS-**--WVRQAPGKGLEWIS*-*----*-*-------RF
Non-T   ................................NYSMN..............SISGSSRYIYYADFVKG..
D10-A   ................................NSGMN..............SIGGSSRYIYYADFVKG..
G5-A    ................................NSGMN..............SIGGSSRYIYYADFVKG..
C1-A    ................................NYYMN..............SISGSSRGIYYADFVKG..
E9-A    ................................NYYMN..............GISGSSRYIYYADFVKG..
B11-A   ................................NYYMN..............GISGSSRYIYYADFVKG..
F6-B    ................................NYYMN..............YISGSSRGIYYADFVKG..
A12-AB  ................................NYGMN..............YISGSSRYIGYADFVKG..
           VH                           CDR1                     CDR2

                80        90       100       110       120       130       140
        ----:----|----:----|----:----|----:----|----:----|----:----|----:----|
fwk     TISRDNAKNSLYLQMNSLRAEDTAVYYCVR-*********----WGRGTLVTVSS---------------
Non-T   .......T......................SSITIFGG--GMDV...........GGGGSGGGGSGGGGS
D10-A   ..............................SNY-------GMDV...........GGGGSGGGGSGGGGS
G5-A    ..............................SDY-------GMDV...........GGGGSGGGGSGGGGS
C1-A    ..............................SSGYSFGY--GMDV...........GGGGSGGGGSGGGGS
E9-A    ..............................SSGYSFGY--GMDV...........GGGGSGGGGSGGGGS
B11-A   ..............................SSGYSFGY--GMDV...........GGGGSGGGGSGGGGS
F6-B    ..............................SSYYSG----GMDV...........GGGGSGGGGSGGGGS
A12-AB  ..............................SSYYY-----GMDV...........GGGGSGGGGSGGGGS
                                   CDR3                  linker

               150       160       170       180       190       200       210
        ----:----|----:----|----:----|----:----|----:----|----:----|----:----|
fwk     QSVLTQPASVSGSPGQSITISC---------***--WYQQHPGKAPKLMIY*--*---GVSNRFSGSKSG
Non-T   ......................AGTSSDVGGYNYVS...............EDSKRPS............
D10-A   ......................AGTSSDVGGSYSVS...............YDSYRPS............
G5-A    ......................AGTSSDVGGDYSVS...............YDSYRPS............
C1-A    ......................AGTSSDVGGGGYVS...............YDSYRPS............
E9-A    ......................AGTSSDVGGNSYVS...............YDSYRPS............
B11-A   ......................AGTSSDVGGSNYVS...............YDSYRPS............
F6-B    ......................AGTSSDVGGYGGVS...............DDSDRPS............
A12-AB  ......................AGTSSDVGGSNYVS...............GDSNRPS............
          VL                          CDR1                  CDR2

               220       230       240       250
        ----:----|----:----|----:----|----:----|
fwk     NTASLTISGLQAEDEADYYC--*-**----FGGGTKLAVL
Non-T   ....................SSYTTRSTRV..........
D10-A   ....................SSSTSYSTRV..........
G5-1    ....................SSGTSYSTRV..........
C1-A    ....................SSGTQGSTRV..........
E9-A    ....................SSGTYYSTRV..........
B11-A   ....................SSGTYGSTRV..........
F6-B    ....................SSGTQYSTRV..........
A12-AB  ....................SSDTGQSTRV..........
                               CDR3
```

**Supplementary Figure S1**. Sequences of the characterized scFv clones. Fwk: conserved framework sequence; Randomized positions in the naive library are indicated with a *. The VH and VL domains are highlighted in green and yellow, respectively. The six CDRs, conforming to Kabat's definition, are highlighted in blue. In the lentiviral constructs, the C-terminal end of the scFv is fused to eGFP via a GAAA linker. Non-T: non-targeted scFv recognizing E. coli beta-galactosidase.

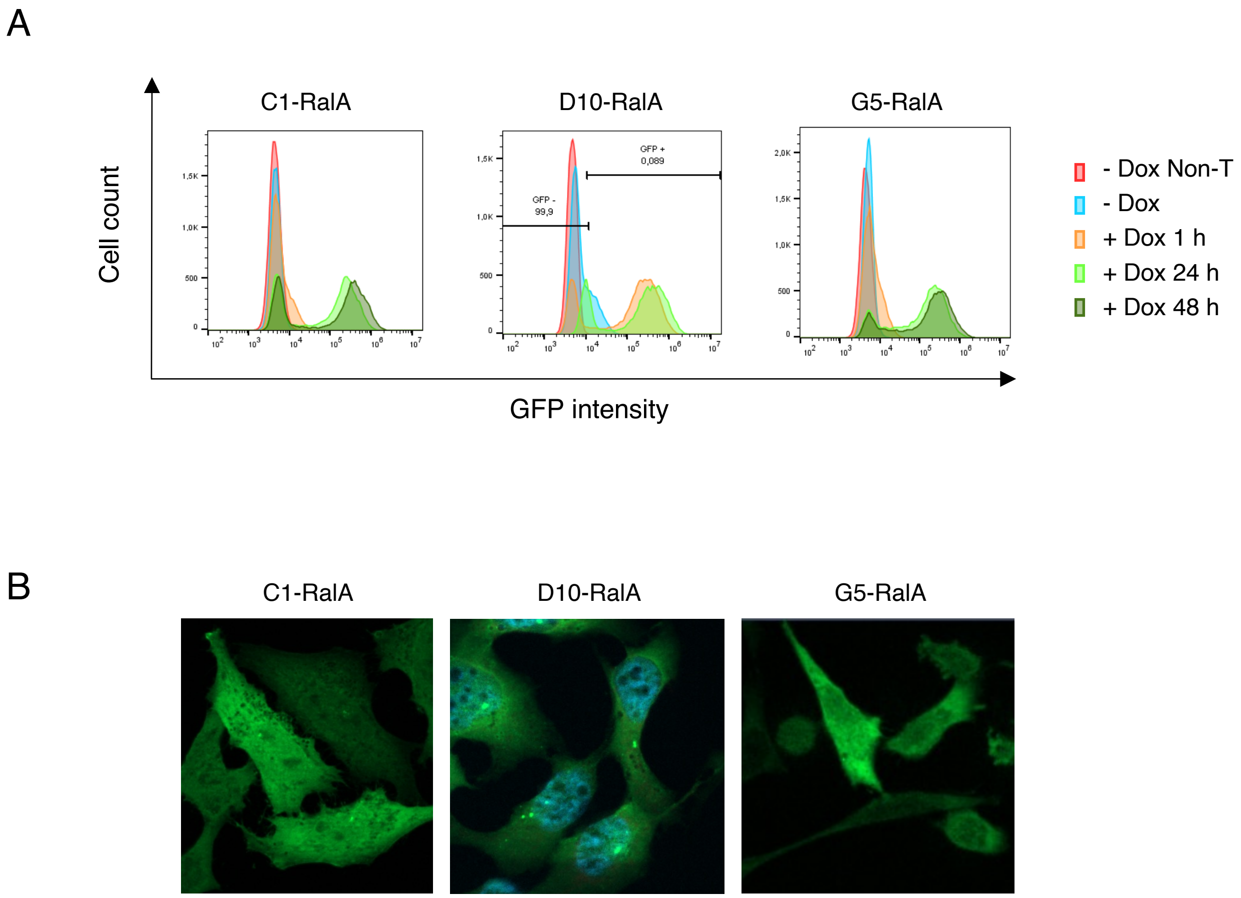


**Supplementary Figure S2.** Expression of anti-RalA clones as intrabodies in HeLa cells. HeLa cells were transduced using lentiviruses expressing scFv-GFP fusions under a dox-inducible promoter. A) Cytometry analysis of the GFP signal. Non-T is the non-targeting intrabody against beta-galactosidase, used as a control (Supplementary Fig. S1). B) The same clones were analyzed at 24 h post-induction by fluorescent microscopy (GFP signal) using a Zeiss Apotome microscope.

A

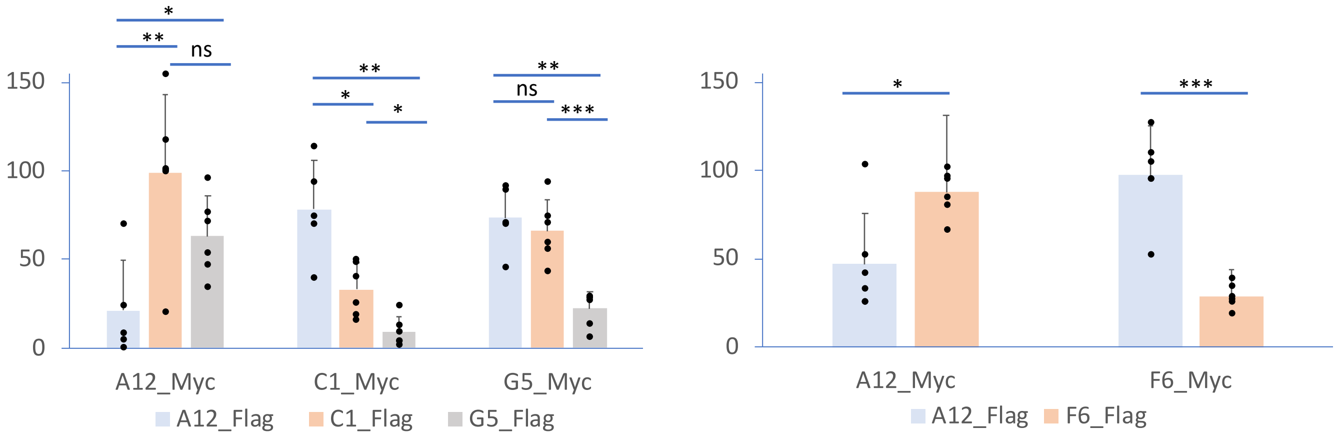


B

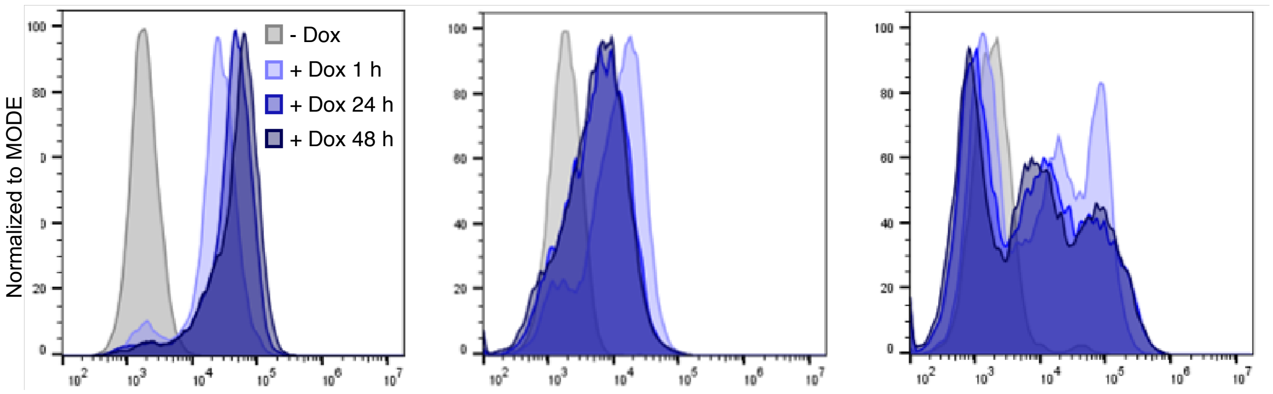


**Supplementary Figure S3.** A) Left: Competition between the 3 anti-RalA clones. Right: Competition between the 2 anti-RalB clones. Competition was determined by ELISA on coated RalA (Left) and RalB (Right). An excess of Myc-tagged antibody was added to the well (30 µg/mL), then a detection was done with a Flag-tagged antibody at a concentration giving a signal proportional to the concentration (2 µg/mL). B) Expression of anti-RalA clones as intrabodies in 4T1 cells. 4T1 cells were transduced using lentiviruses expressing scFv-GFP fusions under a dox-inducible promoter. Cytometry analysis of the GFP signal after 1, 24 and 48h induction by doxycycline.

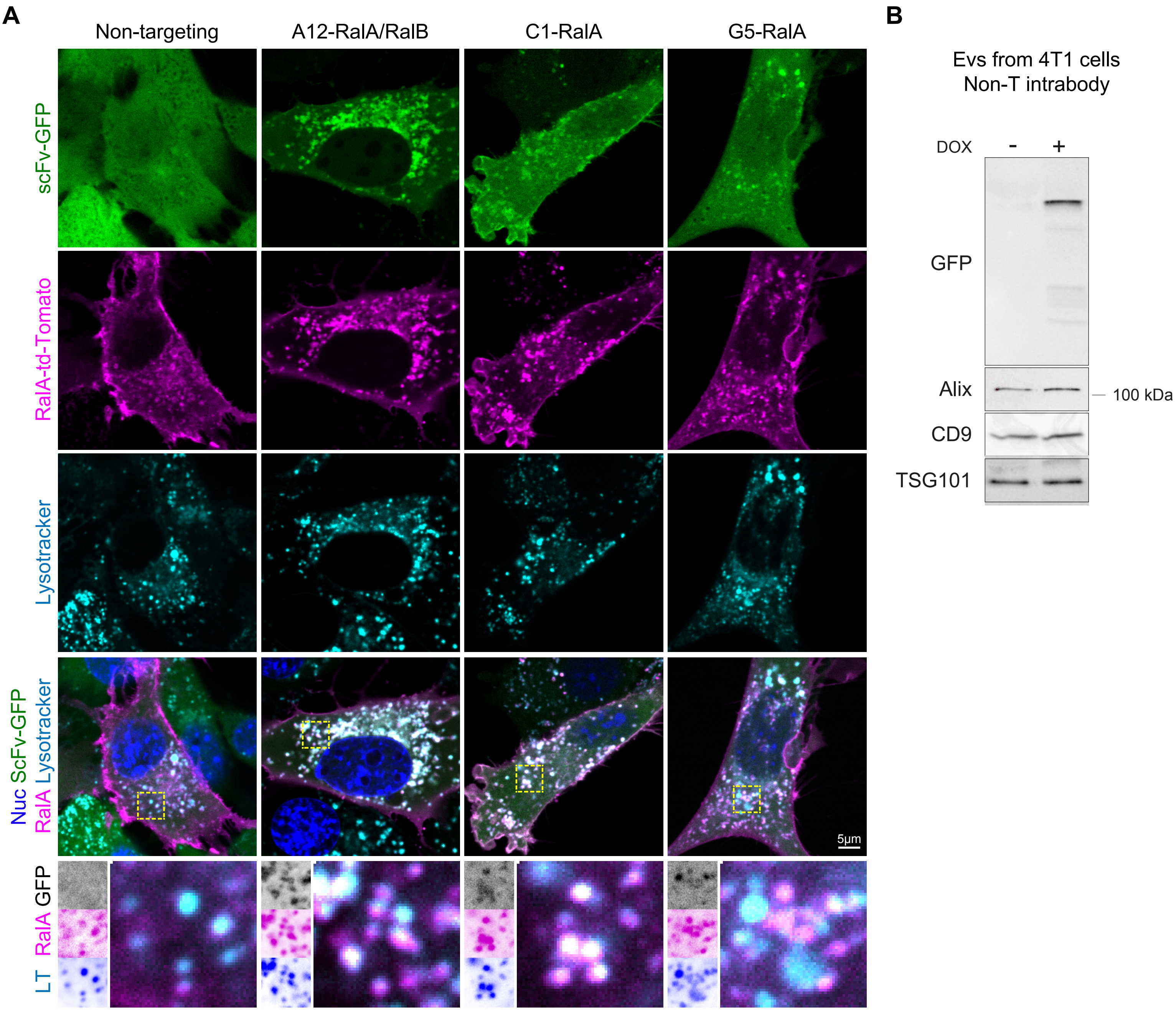


**Supplementary Figure S4**. A) Representative confocal images of 4T1 cells transfected with tdTomato-RalA, stably expressing GFP-coupled intrabodies. Live-cell imaging by Spinning-disk confocal microscopy at 24 h after doxycycline treatment. Scale bar = 5 µm. Intensity adjusted to best visualize the localization patterns. B) EV content is not modified by intrabody expression. Non-T intrabody expression was induced by a 24 h treatment with doxycycline and EVs purified from cell supernatant. EVs (same number per condition) were analyzed by WB using an anti-GFP to detect intrabody expression, and 3 markers of EV, Alix, CD9 and TSG101.
